# Exploring the relationship between *GBA1* host genotype and gut microbiome in the *GBA1*^L444P/WT^ mouse model: Implications for Parkinson disease pathogenesis

**DOI:** 10.1101/2024.12.15.627490

**Authors:** Elisa Menozzi, Mallia Geiger, Victoria Meslier, Federico Fierli, Marine Gilles, Kai-Yin Chau, Aymeric David, Revi Shahar Golan, Alexandre Famechon, Sofia Koletsi, Christian Morabito, Benoit Quinquis, Nicolas Pons, Stanislav Dusko Ehrlich, Jane Macnaughtan, Mathieu Almeida, Anthony HV Schapira

## Abstract

**Background:** Heterozygous variants in *GBA1* are the commonest genetic risk factor for Parkinson disease (PD) but penetrance is incomplete. *GBA1* dysfunction can cause gastrointestinal disturbances and microbiome changes in preclinical models. Mounting evidence suggests that the microbiota-gut-brain axis is potentially implicated in PD pathogenesis. Whether the gut microbiome composition is influenced by host *GBA1* genetics in heterozygosis has never been explored.

**Objectives:** To evaluate whether heterozygosity for the *GBA1* pathogenic L444P variant can cause perturbations in gut microbiome composition.

**Methods:** Faecal samples collected from *GBA1*^L444P/WT^ and *GBA1*^WT/WT^ mice at 3 and 6 months of age were analysed through shotgun metagenomic sequencing.

**Results:** No differences in α- and β-diversity were detected between genotyped groups, at either time points. Overall, we found a little variation of the gut microbiome composition and functional potential between *GBA1*^L444P/WT^ and *GBA1*^WT/WT^ mice over time.

**Conclusion:** Host *GBA1* genotype does not impact gut microbiome structure and composition in the presented *GBA1*^L444P/WT^ mouse model. Studies investigating the effect of a second hit on gut physiology and microbiome composition could explain the partial penetrance of *GBA1* variants in PD.

## Introduction

The aetiology of Parkinson disease (PD) is complex and multifactorial, resulting from both genetic and non-genetic factors (Tanner and Ostrem, 2024). Variants in the *GBA1* gene are found in approximately 10-15% of PD patients, thus representing the commonest genetic risk factor for PD (Petrucci et al., 2020, Skrahina et al., 2021). The *GBA1* gene encodes the lysosomal enzyme glucocerebrosidase (GCase), which breaks down glucosylceramide (GluCer) to ceramide and glucose. Biallelic pathogenic variants in the *GBA1* gene cause Gaucher disease, which is characterised by deficiency in GCase enzymatic activity, resulting in the excessive accumulation of GluCer in multiple innate and adaptive immune cells in the spleen, liver, lung and bone marrow (Pandey et al., 2017).

People with PD carrying *GBA1* variants, especially pathogenic severe variants such as p.L483P (more commonly known as L444P), display a more severe clinical phenotype, characterised by a higher burden of autonomic symptoms and olfactory dysfunction (Carandina et al., 2022, Menozzi and Schapira, 2021). Moreover, severe *GBA1* variants are associated with the highest risk of developing PD (odds ratio up to 30.4) (Vieira et al., 2024). Despite the high frequency of *GBA1* variants detected in PD cases, most individuals carrying homozygous or heterozygous variants in the *GBA1* gene will not develop PD over their lifetime (Hertz et al., 2024). The additional factors that can contribute to increase the risk of developing PD in *GBA1* variant carriers have not yet been identified (Menozzi et al., 2023).

Accumulating research points towards the gastrointestinal tract as a potential initial site of PD pathological changes. Preclinical evidence showed that the caudo-rostral spread of α-synuclein pathology from the gastrointestinal tract to the brain can occur after intestinal inoculation of α-synuclein pre-formed fibrils or α-synuclein derived from human PD brain lysate (Kim et al., 2019, Challis et al., 2020, Holmqvist et al., 2014). Moreover, postmortem and multi-modal imaging studies support the hypothesis that there is a subgroup of patients with PD who could manifest initial α-synuclein pathology in the gastrointestinal tract, namely in the enteric nervous system, with subsequent propagation to the dorsal motor nucleus of the vagus, sympathetic nervous system, brainstem, and the rest of the central nervous system (the so-called ‘body-first’ PD subtype) (Horsager et al., 2020, Borghammer et al., 2022). In contrast, the ‘brain-first’ subtype of PD is characterised by initial deposition of α-synuclein pathology in the olfactory bulb and/or limbic system, with subsequent spread to the brainstem and peripheral nervous system (Horsager et al., 2020). Clinically, body-first and brain-first PD patients would differ, with the former presenting more severe autonomic and olfactory dysfunction compared to the latter (Horsager et al., 2022).

Considering the importance of the gastrointestinal tract in body-first PD pathogenesis, in recent years increasing attention has been drawn to the potential contribution of the gut microbiome in this process. Exposure to gut microbial components such as lipopolysaccharide and lipopeptide enhanced intracellular levels of α-synuclein protein in a murine model of gut enteroendocrine cells (Hurley et al., 2023). Transplantation of faecal material from people with PD promoted motor symptoms and pathological changes in different mouse models, suggesting that alterations in the gut microbiome might trigger PD pathology (Sampson et al., 2016, Sun et al., 2018). Indeed, perturbations of the gut microbiome composition have extensively been reported in several populations of PD patients, without a specific genetic background (Nishiwaki et al., 2024). These perturbations seem to be particularly evident in body-first PD patients compared to brain-first PD (Park et al., 2024).

Because of the similarities in clinical profile displayed by PD patients with pathogenic *GBA1* variants and body-first PD patients, it has been recently hypothesised that *GBA1*-associated PD would more often fit within the body-first PD phenotype (Horsager et al., 2022). Hence, it is reasonable to interrogate the role of the gut microbiome in *GBA1*-associated PD. If and how pathogenic *GBA1* variants *per se* impact on gastrointestinal function and microbiome composition and thus contribute to PD pathogenesis, has been investigated in a very few studies.

Animal models have shown that GCase deficiency can contribute to α-synuclein accumulation in the gastrointestinal tract (Challis et al., 2020). Moreover, heterozygosity for the severe pathogenic L444P *GBA1* variant reduced α-synuclein degradation, induced earlier onset of pathological phosphorylated α-synuclein and exacerbated both motor and gastrointestinal dysfunction in mice also carrying the mutated human *SNCA* A53T (hSNCA^A53T^/*GBA1*^L444P^) compared to hSNCA^A53T^ mice, wild-type for *GBA1* (Fishbein et al., 2014). Delayed intestinal transit time, decreased faecal pellet output and compromised intestinal barrier integrity were detected in aged flies lacking the *Gba1b* gene, the main fly orthologue of *GBA1* (Atilano et al., 2023). In terms of gut microbiome composition, an increased bacterial load and an increased relative abundance of specific genera such as *Acetobacter* and *Lactobacillus* characterised *Gba1b*^*-/-*^ flies; when raised under germ-free conditions, flies showed increased lifespan, improved locomotor abilities, and reduced glial activation (Atilano et al., 2023).

In this study, we aimed to explore for the first time the potential impact of mammalian host severe pathogenic variants in the *GBA1* gene (L444P) on the gut microbiome composition compared to wild-type controls.

## Methods

### Animals

Mice were treated in accordance with local ethical committee guidelines and the UK Animals (Scientific Procedures) Act of 1986. All procedures were carried out in accordance with Home Office guidelines (United Kingdom; Project Licence Number: PP9638474). Male B6;129S4-Gbatm1Rlp/Mmnc (000117-UNC) mice expressing heterozygous knock-in L444P (also known as p.L483P) mutation in the murine *GBA1* gene (*GBA1*^L444P/WT^) were originally purchased from the Mutant Mouse Regional Resource Centre (MMRRC). Littermates (*GBA1*^WT/WT^) were used as controls. Within the first 21 days of life, mice were genotyped as previously described (Migdalska-Richards et al., 2016). After weaning at day 21, they were separated according to genotype into different cages and co-housed together for a maximum of 2 animals per cage. Faecal samples were collected for three consecutive days. On each day of collection, mice were removed from their home cages and singly housed for a maximum of 30 minutes, with water and diet provided during this time. After 30 minutes, faecal samples were collected avoiding urine contamination and immediately placed in airtight Eppendorf tubes into dry ice, until transfer to the laboratory where samples were stored at -80°C prior to further analyses. Faecal samples were collected at 3 and 6 months of age. Faecal pellet output was recorded at each day of collection, and total faecal output was calculated at the end of collection by summing the weights of faecal pellets collected at each day. Animals’ weight was recorded at each collection time point.

### Metagenomics

DNA extraction, high throughput sequencing, read mapping and bioinformatical analysis to determine Metagenomic Species Pangenome (MSP) were performed for the study of the mice microbiota (DOI: dx.doi.org/10.17504/protocols.io.bp2l6×5wklqe/v1 – available after publication).

#### DNA extraction and high throughput sequencing

Frozen faecal materials were aliquoted to ≤250mg and DNA extraction was performed following the procedure previously described in dx.doi.org/10.17504/protocols.io.dm6gpjm11gzp/v1, with the following modifications. 250 μL guanidium thiocyanate, 40 μL N-lauroyl sarcosine (10 % solution) and 500 μL N-lauroyl sarcosine (5 % solution in PBS 1X) were added to each frozen mouse faecal sample, which was then homogenised with a toothpick, vortexed and transferred to a deep-well plate containing 400 μL of 0.1 mm glass beads (not in suspension). Subsequently, the sample plate was incubated at 70°C in a thermomixer for one hour, with stirring at 1,400 rpm. Following centrifugation of the plate at 3,486 ×g for a period of five minutes, the lysate was collected in a new plate. The pellet was then washed with 500 μL of TENP (50 mM Tris-HCL 20 mM EDTA 10 mM NaCl, saturated with PVPP). The plate was vortexed and centrifuged at 3,486 ×g for five minutes, after which the recovered lysate was pooled with the previous one. Finally, the final lysate was centrifuged for 10 minutes at 3,486×g, after which 800 μL were collected in a new plate. This plate was employed for purification with magnetic beads on the QIASymphony. The utilised protocol has been designed for MGP with the QIAGEN DSP Virus/Pathogen kit. DNA was quantified using Qubit Fluorometric Quantitation (ThermoFisher Scientific, Waltham, US) and qualified using DNA size profiling on a Fragment Analyzer (Agilent Technologies, Santa Clara, US). One μg of high molecular weight DNA (>10 kbp) was used to build the library. Shearing of DNA into fragments of approximately 150 bp was performed using an ultrasonicator (Covaris, Woburn, US) and DNA fragment library construction was performed using the Ion Plus Fragment Library and Ion Xpress Barcode Adapters Kits (ThermoFisher Scientific, Waltham, US). Purified and amplified DNA fragment libraries were sequenced using the Ion Proton Sequencer (ThermoFisher Scientific, Waltham, US), with a minimum of 20 million high-quality 150 bp reads generated per library (Meslier et al., 2022).

#### Read Mapping

Reads were quality filtered to remove any low-quality sequences using Alientrimmer software (Criscuolo and Brisse, 2013) and potential host-related reads using Bowtie2 (Langmead and Salzberg, 2012). Resulting high-quality reads were mapped onto the 5 million gene integrated reference catalogue of the Murine Intestinal Microbiota Integrated Catalog v2 (MIMIC2) (Plaza Onate et al., 2021) using the METEOR software suite (Pons et al., 2010). Read mapping was performed in a two-step procedure, using an identity threshold of 95% to the reference gene catalogue with Bowtie2 (Langmead and Salzberg, 2012). First, unique mapped reads were attributed to their corresponding genes. Second, shared reads were weighted according to the ratio of unique mapping counts. A downsizing procedure was performed to normalize the gene counts between samples by randomly selecting a subset of reads depending on the sequencing depth (usually ≥10M reads for an average 20M reads depth sequencing). The gene abundance table was then normalized using the FPKM strategy and analysed using MetaOMineR (momr) R package (https://cran.r-project.org/web/packages/momr/index.html) (Le Chatelier and Prifti).

#### MSP microbial species determination

MSPs were used to quantify species associated to the million gene MIMIC2 reference catalogue (Nielsen et al., 2014, Plaza Onate et al., 2019). MSPs are clusters of co-abundant genes (min size ≥ 500 genes) that likely belong to the same microbial species, reconstructed from the 5 million genes catalogue into 1252 MSPs. MSP abundance profiles were calculated as the mean abundance of 100 markers genes, defined as the robust centroids of each MSP cluster. A threshold of 10% of the marker genes was applied as MSP detection limit. Taxonomical annotation was performed using GDTB R07-RS207 (Parks et al., 2022).

#### Assessment of the microbial functional potentials

To determine functional potential of the gut microbiota at the module level, we used an INRAE pipeline as previously described (Thirion et al., 2023). Three databases were used to predict gene functions: Kyoto Encyclopedia of Genes and Genomes (KEGG) (Kanehisa and Goto, 2000), eggNOG database (version 3.0) (Huerta-Cepas et al., 2016), and TIGRFAMs (version 15.0) (Haft et al., 2001). First, genes of the 5 million genes catalogue were annotated using KEGG107 database using Diamond (Buchfink et al., 2015) and further clustered into functional pathway modules according to KEGG Orthology (KO) groups, Gut Metabolic Modules (GMM) (Vieira-Silva et al., 2016), and Gut Brain Modules (GBM) (Valles-Colomer et al., 2019). Second, KEGG, GMM and GBM modules were reconstructed in each MSP using their reaction pathways based on their detected annotated KO, NOGs or TIGRFAMs genes. GMM and GBM functional modules were selected because they are specific to gut bacterial and gut-brain axis functions. For each pair of MSP/mouse, the completeness of any given functional modules was calculated by considering the set of genes detected in the MSP of each mouse and the MSP completeness in each mouse. For a given MSP in a specific mouse, completeness of the modules was corrected by the abundance of the MSP. After correction, functional modules in each MSP/mouse were considered as complete if at least 90% of the involved reactions were detected. Abundance of functional modules in each sample was computed as the sum of the MSP abundances containing the complete functional module.

### Statistical analysis

Statistical analysis and visualisation were performed with R software (version 4.4.1) (R-Core-Team, 2019).

Group differences (WT/WT vs L444P/WT) in faecal pellet output, animal weights, richness, α□diversity measures, and contrasts in species and functional modules abundances were computed using non-parametric Wilcoxon rank-sum test. For comparisons over time, the Wilcoxon signed-rank test was applied. Effect size was calculated as Cliff’s Delta. Differences in the prevalence of each species in each group, defined as the proportion of samples containing that species in relation to the total number of samples in the respective group, were calculated using Fisher’s exact test. Shannon diversity index was computed based on the MSP matrix using the function *diversity* from the R package vegan (Dixon, 2003). Principal coordinates analysis (PCoA) was performed on the Bray-Curtis dissimilarity index with the R package ade4 (Thioulouse et al., 2018). Permutational multivariate analysis of variance (PERMANOVA) was computed using distance matrices with the function *adonis2* from the R package vegan (n = 1000 permutations), to assess differences in β□diversity (Dixon, 2003).

## Results

### Study cohort

An overview of the characteristics of the study cohort is presented in Table 1. A total of 16 WT/WT and 13 L444P/WT animals were included in the study (with 1 L444P/WT sampled only at 3 months). No difference in the total faecal output weight was detected at 3 or 6 months between genotypes, or between time points within each group. We observed a significant weight gain between 3 and 6 months of age within the WT/WT group (p= 0.000031) and within the L444P/WT group (p=0.00049). At 6 months, the median weight of the L444P/WT group was significantly higher than the one in the WT/WT group (p = 0.017), but the difference between 3 and 6 months (delta weight) only showed a trend towards significance, being higher in L444P/WT (p=0.053).

**Table 1.** Overview of study cohort.

| **Genotype** | **WT/WT** |  | **L444P/WT** |  | ***p value*** |  |
| --- | --- | --- | --- | --- | --- | --- |
| **Age (months)** | 3 | 6 | 3 | 6 | P1 | P2 |
| **N** | 16 | 16 | 13 | 12 |  |  |
| **Faecal output (g)** | 0.72 | 0.67 | 0.62 | 0.78 | ns | ns |
| **Animal weight (g)** | 31.50 | 36.67 | 33.35 | 42.06 | ns | 0.017 |

Legend: data are presented as median values. P1 = comparison between WT/WT and L444P/WT at 3 months; P2 = comparison between WT/WT and L444P/WT at 6 months. P values, as determined by Wilcoxon rank-sum test, are reported in the table (ns = not significant).

### Limited variation of the gut microbiome structure between groups

Gene count and MSP richness were not found to be significantly different between groups at different time points (p=0.33 and p=0.55 for MSP richness, at 3 and 6 months, respectively, Fig. 1A). This finding was further confirmed by analysing additional α-diversity indices, including the Shannon index. The Shannon index was 3.61 and 3.67 for the WT/WT and L444P/WT groups at 3 months (p=0.47), and 3.61 and 3.71 at 6 months (p=0.66), respectively. In line with weight, MSP richness was found to increase over time in each group (p=0.012 and p=0.034 for MSP richness in the WT/WT and L444P/WT groups, respectively, Fig. 1B).

**Figure 1.**
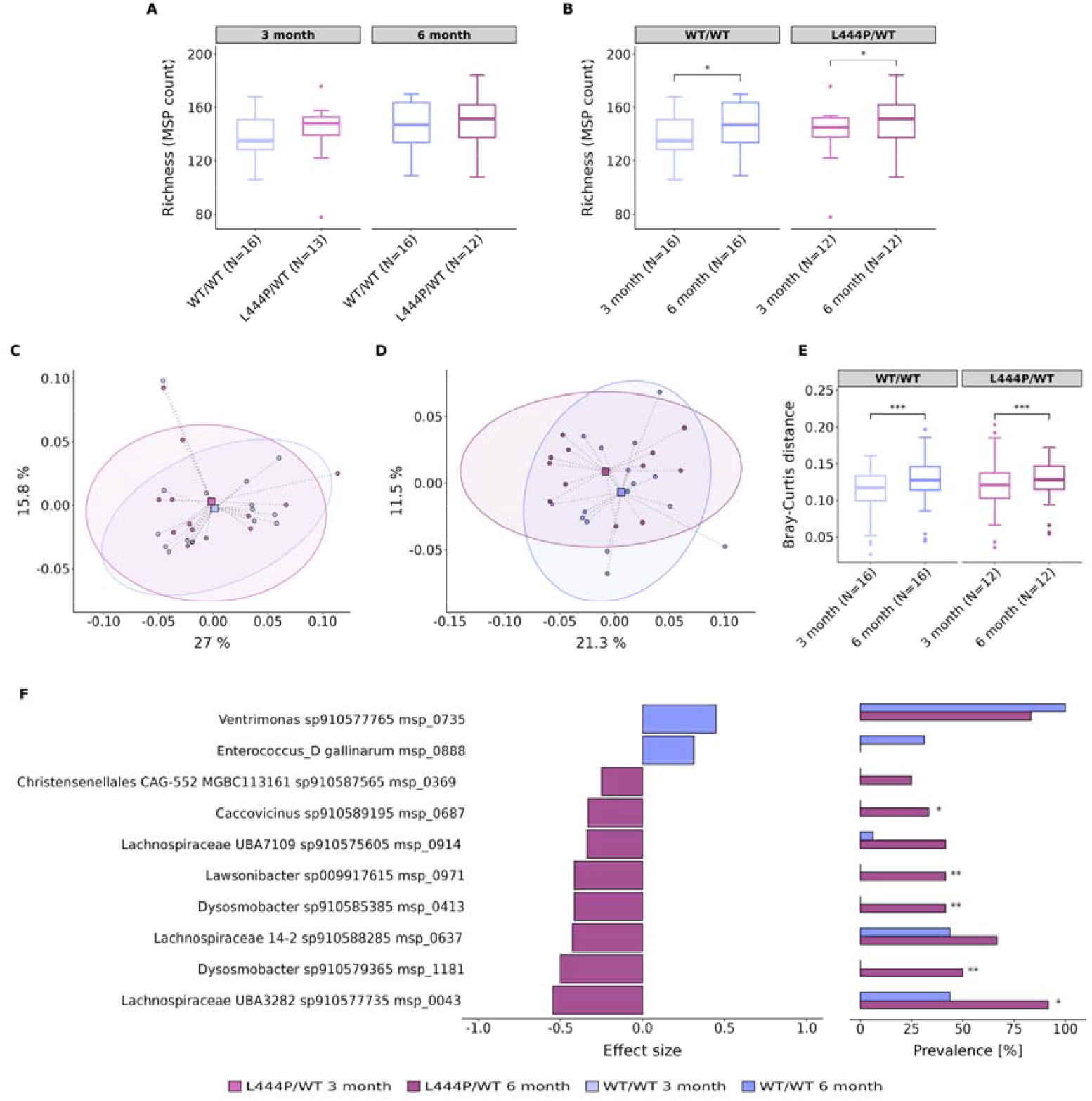
Gut microbiome alterations in *GBA1*^L444P/WT^ and *GBA1*^WT/WT^ mice. (A) Differences in species (MSP) richness between groups (WT/WT and L444P/WT) at 3 months and 6 months. (B) Differences in species (MSP) richness within the WT/WT group and the L444P/WT group over time. PCoA performed on the Bray-Curtis dissimilarity index at 3 months (C) and 6 months (D). (E) Differences in Bray-Curtis dissimilarity index within the WT/WT and L444P/WT groups at different times points. (F) Microbial species abundance between L444P/WT and WT/WT animals at 6 months: left, diverging bar plots show the relative abundance of species in WT/WT and L444P/WT according to the effect size measured by the Cliff’s Delta; right, bar charts show the percentage of prevalence of species between groups (calculated as number of samples presenting the species compared to the total number of samples in each group). Significant differences, determined using Wilcoxon’s signed-rank test and Fisher’s exact test, are indicated by a star above each box plot and bar plot (* = p<0.05, ** = p<0.01).

We also performed β-diversity analysis as measured by the Bray-Curtis dissimilarity index. We did not observe a significant difference between genotypes at either of the time points (p=0.73 at 3 months and p=0.52 at 6 months, Fig. 1C and 1D respectively), although a significant variation of the structure of the gut microbiome between 3 and 6 months of age was identified in both groups (p=0.0009 for the WT/WT, and p=0.0009 for the L444P/WT, Fig. 1E).

### Differentially abundant species and functional modules between groups

At 3 months, only one species was differentially abundant between groups, being enriched in the L444P/WT group (*Schaedlerella sp910575475*), confirming the little variation of the gut microbiome observed at 3 months of age. At 6 months, there were 10 species differentially abundant, with a majority of them only detected in one group (Fig. 1F). Two species, detected in both groups but more abundant in the L444P/WT, belonged to the family *Lachnospiraceae*, phylum Firmicutes_A. At 6 months, the proportion of enriched species in the L444P/WT group was higher than in the WT/WT group (Fisher’s exact test: p=0.02).

In line with the limited differences observed for the microbial species, little variation was found for the functional potential between groups at each time point. Only one KEGG module enriched in WT/WT (ascorbate degradation, ascorbate => D-xylulose-5P, M00550) and one GMM module enriched in L444P/WT (glycocholate degradation, MF0044) were found to be differentially abundant between groups at 3 months. At 6 months, the same GMM module enriched at 3 months in the L444P/WT (MF0044) was also enriched, whereas no differences in KEGG or GBM modules were found between genotypes.

## Discussion

To the best of our knowledge, this is the first study to investigate the gut microbiome composition of mice carrying *GBA1* variants in heterozygosis, the most frequent genetic risk factor for PD. Our data did not show any significant perturbations of the gut microbiome structure and abundance of species or functional modules due to standalone *GBA1* genetic status.

These findings are in line with the available literature on Gaucher disease. Gastrointestinal involvement is very rarely reported in Gaucher disease and when present, is largely unresponsive to enzyme replacement therapy (Kim et al., 2017). Perturbations of gut microbiome leading to fat malabsorption and intestinal dysfunction have been described in knockout mice for the *SCARB2* gene, which encodes the lysosomal integral membrane protein-2 (LIMP-2) which is fundamental to traffic GCase to the lysosome (Li et al., 2024), however no report on gut microbiome in patients with Gaucher disease is currently available.

*GBA1* variant carriers represent a group of susceptible individuals at-risk of future development of PD, but the limited penetrance of *GBA1* variants for PD implies that additional factors are needed for clinically manifest disease expression. It has been suggested that in susceptible individuals, exposure to ingested toxicants or intestinal infections could result in low-grade gut inflammation and microbiome alterations, which could instigate α-synuclein pathology in the gut with subsequent rostral spread to the brain (Houser and Tansey, 2017, Dorsey et al., 2024, Khare et al., 2019, Wang et al., 2021, Ilieva et al., 2022, Zeng et al., 2022). Previous models of mild chronic colitis using dextran sodium sulfate (DSS) induced more severe motor dysfunction, microglia activation, and dopaminergic neuron loss in genetic models of PD such as mice carrying the *LRRK2* G2019S variant (Lin et al., 2022). Recent studies investigating the effect of DSS or paraquat, a neurotoxic herbicide, found that these stimuli differentially interacted with specific genotypes including the hSNCA^A53T^/hSNCA^A53T^ or the hSNCA^A53T^/*GBA1*^L444P^, leading to toxicant-genotype specific microbiome alterations, gastrointestinal dysfunction and neurodegenerative changes (Chaklai et al., 2024). We could therefore hypothesise that the combination of *GBA1* host genetics and exposure to ingested toxicants and/or gastrointestinal inflammation (dual-hit hypothesis) could represent a possible pathogenetic mechanism underlying *GBA1*-PD development which is mediated by gut microbiome perturbations and gastrointestinal dysfunction.

We acknowledge that our study has some limitations. First, mice were sacrificed at 6 months of age, so potential changes in the gut microbiome consequent to ageing might have been missed, although in our pilot study we did not observe any dramatic difference in gut microbiome composition between 6 months and later time points (data shown in Supplementary Material). Second, we did not perform any measurement of gastrointestinal transit time, inflammation or permeability.

Notwithstanding, the design of our study (different genotypes kept in separate cages within the same biological facility and under the same dietary conditions) minimised the risk of gut microbiome homogenisation between different genotyped animals due to coprophagia, as previously observed (Supplementary Figure 1). Thus, our study represents an essential preliminary step to evaluate the validity of the dual-hit hypothesis for *GBA1*-PD pathogenesis in future models.

## Acknowledgment

We would like to thank the UCL Royal Free BSU staff for their precious help with the study execution.

This research was funded in part by Aligning Science Across Parkinson’s (Grant number: ASAP-000420) through the Michael J. Fox Foundation for Parkinson’s Research (MJFF) and by the EU Joint Programme—Neurodegenerative Research (JPND) through the MRC grant code MR/T046007/1. Additional funding was provided from the MetaGenoPolis grant ANR-11-DPBS-0001. For the purpose of open access, the author has applied a CC BY 4.0 public copyright license to all Author Accepted Manuscripts arising from this submission.

## Authors’ Roles

1. Research project: A. Conception, B. Organization, C. Execution.
2. Statistical Analysis: A. Design, B. Execution, C. Review and Critique.
3. Manuscript Preparation: A. Writing of the first draft, B. Review and Critique.

EM: 1A, 1B, 1C, 2A, 2B, 2C, 3A.

MGe: 2B, 2C, 3A, 3B.

VM: 1A, 2A, 2B, 2C, 3A, 3B.

FF: 1B, 1C, 3B.

DC: 1A, 1B, 3B.

RSG: 1B, 1C, 3B.

SK: 1B, 3B.

CM: 1C, 3B.

AD: 1C, 3B.

MGi: 1C, 3B.

BQ: 1C, 3B.

AF: 1C, 3B.

NP: 2B, 3B.

SDE: 1A, 2C, 3B.

JM: 1A, 1B, 3B.

MA: 1A, 1B, 2C, 3B.

AHVS: 1A, 3B.

## Financial Disclosure/Conflict of Interest concerning the research related to the manuscript

The authors report no conflict of interest.

