## Supplementary Material for "Exploring the relationship between *GBA1* host genotype and gut microbiome in the *GBA1*^L444P/WT^ mouse model: Implications for Parkinson disease pathogenesis"

**Table of Content:**

- **Supplementary Methods (p. 2)**
- **Supplementary Results (p. 3-4)**
- **Supplementary Tables (p. 5)**
- **Supplementary Figure 1 (p. 6)**

**Pilot study to evaluate cage and time effect**

**Supplementary Methods**

Study protocol and methodology of this pilot study were similar to the one presented for the main study.

The pilot study differed from the main study in the following aspects. First, the two genotypes (L444P/WT and WT/WT) were co-housed and mixed for a maximum of 5 animals per cage. Second, faecal samples were collected at four consecutive time points (3, 6, 9, and 12 months of age).

**Statistical analysis**

Continuous variables are presented as means ± SDs. Differences between genotyped groups in faecal pellet weights, richness, and abundances of species/functional modules, were computed using t-test or non-parametric Wilcoxon rank-sum test, according to normality of the variables. When appropriate, p-values were corrected for multiple testing with the Benjamini-Hochberg (BH) false discovery rate (FDR) correction, with a significance threshold of FDR-corrected p<0.05.

To test differences of the microbiome variables over time within the WT/WT and L444P/WT groups separately, one-way repeated measures analysis of variance by ranks using Friedman test was applied, with post-hoc analysis with Nemenyi test if Friedman test was significant.

A corrected p value (q value) was considered significant if below 0.2. If unadjusted, p values were considered significant if below 0.05.

**Supplementary Results**

**Study cohort**

Demographics characteristics of the study cohort of the pilot study are shown in Supplementary Table 1. Eight WT/WT and nine L444P/WT mice have been included, with five mice per group completing the 4-time points collection (thus called complete cases). Different genotypes were mixed into five cages as follows:

- cage 1 contained two WT/WT and two L444P/WT mice;
- cage 2 contained two WT/WT and two L444P/WT mice;
- cage 3 contained one WT/WT and one L444P/WT mice;
- cage 4 contained one WT/WT and two L444P/WT mice;
- cage 5 contained two WT/WT and two L444P/WT mice.

The mean total faecal output did not differ between the WT/WT and L444P/WT groups at any of the time points.

**Cage effect**

Principal component analysis (PCoA) ordination of Bray-Curtis dissimilarity index did not show a significant effect of genotype (p adonis = 0.56). We found a significant effect of cage (p adonis = 0.001; Supplementary Fig. 1) and of time, although to a lesser extent (p adonis = 0.047).

**Gut microbiome structure**

Gene count (GC) and MSP richness results are reported in Supplementary Table 2. No differences in GC or MSP species richness were observed between genotypes at 3, 6, 9 and 12 months.

Using only the complete cases and repeated measures analysis on paired samples within each group (WT/WT and L444P/WT), we did not detect any significant difference over time in GC richness in the WT/WT group, χ^2^(3) = 3.48, p = 0.32, or in the L444P/WT group, χ^2^(3) = 2.52, p = 0.47. For MSP richness, there was a statistically significant difference over time in the WT/WT group, with a large effect (W = 0.584). Post-hoc Nemenyi test revealed statistically significant differences between 3 and 12 months (q = 0.017). In the L444P/WT group, we found similar results with a large effect (W = 0.808) and a statistically significant difference in MSP species richness between 3 and 12 months (q = 0.0034).

**Abundance of contrasted microbial species**

No significant differences in species abundance were detected between WT/WT and L444P/WT mice, after correction for multiple testing. A few contrasted species were identified at each time point when we considered unadjusted comparisons (5 species enriched in the WT/WT group at 3 months; 1 species enriched in the WT/WT group at 6 months; 4 species contrasted at 9 months, with 3 enriched in the WT/WT group and 1 in the L444P/WT group; and 5 species contrasted at 12 months of age, with 2 enriched in the WT/WT group and 3 in the L444P/WT group).

Using Friedman test, there were no contrasts in species abundance within WT/WT or L444P/WT groups over time which resisted to correction for multiple testing, suggesting stability of the gut microbiome composition within each group over time. When we considered unadjusted comparisons, only 16 and 13 contrasted MSP species were detected in the WT/WT and L444P/WT groups, respectively, over time.

**Functional modules abundance**

No contrasts in the abundance of functional modules resisted to correction for multiple testing, when WT/WT and L444P/WT groups were compared. A few contrasted functional modules were identified at each time point, when we considered unadjusted comparisons (3 and 2 functional modules were enriched in the WT/WT group at 3 and 6 months; no contrasts at 9 months; 10 modules at 12 months, of which 7 enriched in WT/WT and 3 in L444P/WT).

Using Friedman test, there were no contrasts in functional modules within WT/WT or L444P/WT groups over time which resisted to correction for multiple testing. When we considered unadjusted comparisons, only 11 and 16 modules pathways contrasted over time within the WT/WT and L444P/WT groups, respectively. Thus, the functional potential analysis reflected what observed for the microbial species analysis.

Overall, these findings suggest that A) co-housing animals with different genotypes in the same cage can be dominant to genotype effect in determining gut microbiome diversity and composition, and thus to minimise the “cage effect”, mice with different genotypes should be kept in separate cages, and B) the gut microbiome composition of mice, either WT/WT or L444P/WT, is relatively stable at time points later than 6 months.

**Supplementary Tables**

**Supplementary Table 1. Characteristics of study cohort.**

| **Genotype** | **Time point (months)** | **N** | **Mean total faecal output (weight – g)** |
| --- | --- | --- | --- |
| **WT/WT** | 3 | 8 | 0.47 |
| **WT/WT** | 6 | 8 | 0.5 |
| **WT/WT** | 9 | 7 | 0.62 |
| **WT/WT** | 12 | 5 | 0.55 |
| **L444P/WT** | 3 | 9 | 0.42 |
| **L444P/WT** | 6 | 8 | 0.49 |
| **L444P/WT** | 9 | 6 | 0.76 |
| **L444P/WT** | 12 | 5 | 0.66 |

**Supplementary Table 2. GC and MSP species richness in WT/WT and L444P/WT.**

|  | **Time (months)** | **WT/WT** | **L444P/WT** | ***p value*** | ***q value*** |
| --- | --- | --- | --- | --- | --- |
| **GC richness** |  |  |  |  |  |
|  | 3 | 367440.8 ± 41023.73 | 354683.3 ± 48401.95 | 0.61 | 0.96 |
|  | 6 | 382006.2 ± 43779.72 | 374760.4 ± 23484.72 | 0.72 | 0.96 |
|  | 9 | 373824.4 ± 24326.82 | 371917.3 ± 38364.78 | 0.84 | 0.96 |
|  | 12 | 419525.6 ± 60521.92 | 388214.3 ± 19937.12 | 0.31 | 0.80 |
| **MSP richness** |  |  |  |  |  |
|  | 3 | 116 ± 8.60 | 114.67 ± 8.35 | 0.73 | 0.85 |
|  | 6 | 120.12 ± 8.69 | 122.12 ± 8.25 | 0.6 | 0.73 |
|  | 9 | 120 ± 7.32 | 123.17 ± 1.33 | 0.38 | 0.51 |
|  | 12 | 126 ± 15.71 | 131 ± 3.16 | 1 | 1 |

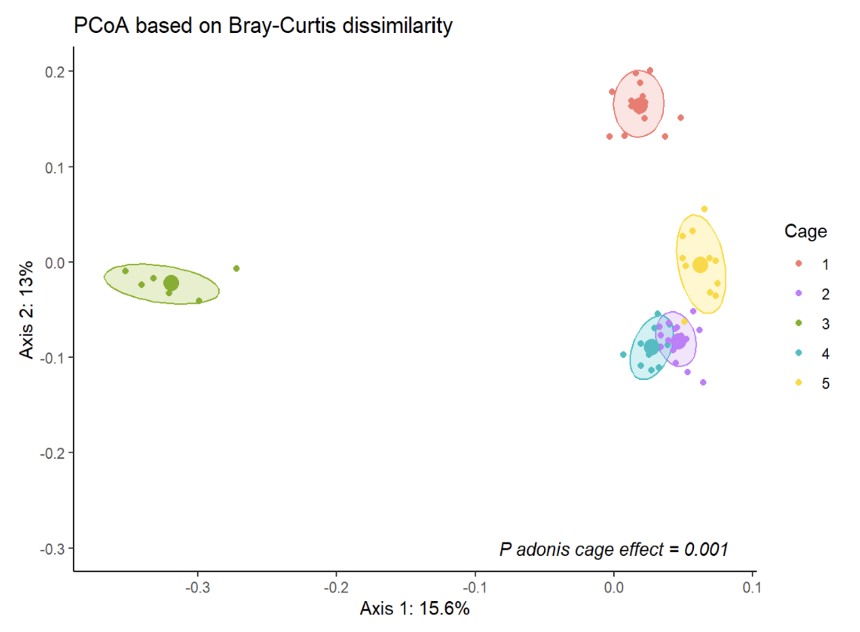

**Supplementary Figure 1. PCoA plots based on Bray-Curtis dissimilarity with PERMANOVA testing the cage effect (n = 999 permutations).**
